# Bacterial Cytological Profiling Identifies Rhodanine-containing PAINS Analogs as Specific Inhibitors of *E. coli* Thymidylate Kinase *in Vivo*

**DOI:** 10.1101/2021.02.22.432404

**Authors:** Elizabeth T. Montaño, Jason F. Nideffer, Joseph Sugie, Eray Enustun, Adam B. Shapiro, Hannah Tsunemoto, Alan I. Derman, Kit Pogliano, Joe Pogliano

## Abstract

In this study, we conducted an activity screen of 31 structural analogs of rhodanine-containing pan-assay interference compounds (PAINS). We identified nine active molecules inhibiting the growth of *E. coli* and classified them according to their *in vivo* mechanisms of action. The mechanisms of action of PAINS are generally difficult to identify due to their promiscuity. However, we leveraged bacterial cytological profiling, a fluorescence microscopy technique, to study these complex mechanisms. Ultimately, we found that although some of our molecules promiscuously inhibit multiple cellular pathways, a few molecules specifically inhibit DNA replication despite their structural similarity to related PAINS. A genetic analysis of resistant mutants revealed that thymidylate kinase (an enzyme essential for DNA synthesis) is an intracellular target of some of these rhodanine-containing antibiotics. This finding was supported by assays of *in vitro* activity as well as experiments utilizing a thymidylate kinase overexpression system. The analog that demonstrated the lowest IC_50_ *in vitro* and MIC *in vivo* displayed the greatest specificity for the inhibition of DNA replication in *E. coli*, despite containing a rhodamine moiety. While it’s generally thought that PAINS cannot be developed as antibiotics, this work highlights the utility of bacterial cytological profiling for studying the *in vivo* specificity of antibiotics, and it showcases novel inhibitors of *E. coli* thymidylate kinase.

**Importance:** We demonstrate that bacterial cytological profiling is a powerful tool for directing antibiotic discovery efforts because it can be used to determine the specificity of an antibiotic’s *in vivo* mechanism of action. By assaying analogs of PAINS, molecules that are notoriously intractable and non-specific, we (surprisingly) identify molecules with specific activity against *E. coli* thymidylate kinase. This suggests that structural modifications to PAINS can confer stronger inhibition by targeting a specific cellular pathway. While *in vitro* inhibition assays are susceptible to false positive results (especially from PAINS), bacterial cytological profiling provides the resolution to identify molecules with specific *in vivo* activity.

## Introduction

Due to the increasing emergence of multi-drug resistant (MDR) bacteria, there is a desperate need for new antibiotics to effectively treat persistent infections.^1–3^ In 2015, ceftazidime-avibactam-resistant *Klebsiella pneumoniae* was identified within months of the antibiotic’s approval for use in the clinic.^4^ And, even as the rate at which pathogens are developing resistance to clinically administered antibiotics continues to increase, many companies once actively involved in antibiotic discovery are abandoning these efforts.^5–11^ Consequently, the speed at which antibiotics are achieving clinical status is not fast enough to keep pace with evolving resistance.^9–11^ For this reason, the continued research and development of potential drug candidates is critical for combatting one of the greatest threats to global health today.

Vast libraries of natural products and/or synthetic molecules are frequently screened in order to identify those with antibacterial activity.^12^ These screens often yield a number of active molecules; however, potent inhibitors are rarely pursued if their mechanisms of action (MOA) are difficult to classify. Consequently, many of these molecules do not make it to the development stage of antibiotic research.^13^ Pan-assay interference compounds (PAINS) constitute a structurally diverse class of molecules that contain highly reactive functional groups (such as toxoflavin, isothiazolone, curcumin, hydroxyphenol hydrazine, phenol-sulfonamide, or rhodanine) that frequently interfere with the integrity of biological assays and yield false positive results.^14^ Their tendency to react with or bind promiscuously to many different targets makes PAINS notoriously difficult to study, and for this reason, they are often discarded from antibiotic investigations.^15, 16^ In fact, filters have been developed for removing PAINS from compound libraries prior to screening.^17^ Consequently, many PAINS remain unstudied.

Previously, we developed bacterial cytological profiling (BCP) as a method for rapidly pinpointing the cellular pathway(s) targeted by an antibiotic.^18, 19^ BCP, which has been used to ascertain the MOA of a variety of compounds active against Gram-negative and Gram-positive bacteria, is predicated on the different cytological changes that occur in cells exposed to antibiotics with various mechanisms.^18–22^ For example, inhibitors that strongly block protein synthesis (i.e. tetracycline) cause the DNA to assume a toroidal structure, while inhibitors of transcription (i.e. rifampicin) cause the DNA to decondense (Figure S1). Inhibitors of cell wall biogenesis induce cell shape defects and cause lysis, while antibiotics affecting DNA replication (i.e. ciprofloxacin) yield elongated cells that contain a single, centrally located nucleoid. We previously demonstrated that the inhibition of multiple pathways within a single cell results in a combination of distinguishable phenotypes.^22, 23^ Thus, BCP enables us to study mechanistically complex molecules, which would otherwise present an insurmountable challenge due to their multiple cellular targets or general reactivity. When used for SAR analyses, BCP facilitates the identification of molecular analogs that have a greater specificity for a single target pathway.

Here, we conducted an antibiotic screen and identified a chemical series of PAINS-related molecules with antibacterial activity against *E. coli ΔtolC*. With the understanding that PAINS have little direct clinical relevance, we pursued these molecules in order to validate BCP as an approach for studying mechanistically complex antibiotics. Using BCP, we discovered that some PAINS-related molecules inhibit multiple pathways *in vivo*, but other structural analogs exhibited high specificity for the DNA synthesis pathway. *In vivo* and *in vitro* experiments revealed that some of these molecules specifically inhibit thymidylate kinase (TMPK), an essential enzyme that catalyzes the synthesis of thymidine 5’-diphosphate from thymidine 5’-monophosphate.^24^ Conserved in many bacterial species, TMPK is a promising target for new antibiotics.^25^ Thus, our study identifies potentially powerful cell biology tools: molecules with specific activity against TMPK in an isolated *E. coli* system as well as a novel application of BCP for studying families of antibiotics that have long been considered too much of a “pain” to study.

## Results and Discussion

### Characterizing the cellular pathways inhibited by two PAINS using BCP and viability studies

A library of 1,798 synthetic small molecules was screened to identify those that inhibit the growth of *E. coli ΔtolC*. In conducting this screen, we identified two active compounds that shared a similar backbone structure containing a rhodanine moiety, suggesting that they might be PAINS^9^. These molecules, designated compound 1 and compound 2, are shown in Table 1. The MICs of compounds 1 and 2 against *E. coli ΔtolC* were determined to be 11.7 *μ*M and 6.1 *μ*M, respectively. To rapidly identify the pathway(s) inhibited by these compounds, we performed BCP. *E. coli ΔtolC* cells were treated with each compound at concentrations equivalent to 1X, 3X, and 5X MIC for durations of 30 minutes, one hour, two hours, and four hours. Following treatment, the cells were stained with fluorescent dyes and imaged using high-resolution fluorescence microscopy (Figure 1). Cells treated with compound 1 at 3X and 5X MIC for two hours or more were elongated and contained a single, centrally located nucleoid (Figure 1A), resembling cells treated with known DNA replication inhibitors (Figure S1).^19–22^ At these concentrations, some cells also appeared to be swollen or lysed, typical of cells treated with cell wall inhibitors. Compound 2 also induced DNA replication defects after one hour of treatment at all tested concentrations, and a severe degree of cell lysis and spheroplast formation were observed after four hours of treatment (Figure 1B). Thus, consistent with the knowledge that PAINS often target multiple pathways, compounds 1 and 2 inhibited two distinct metabolic processes, DNA replication and cell wall biogenesis.

**Table 1.**
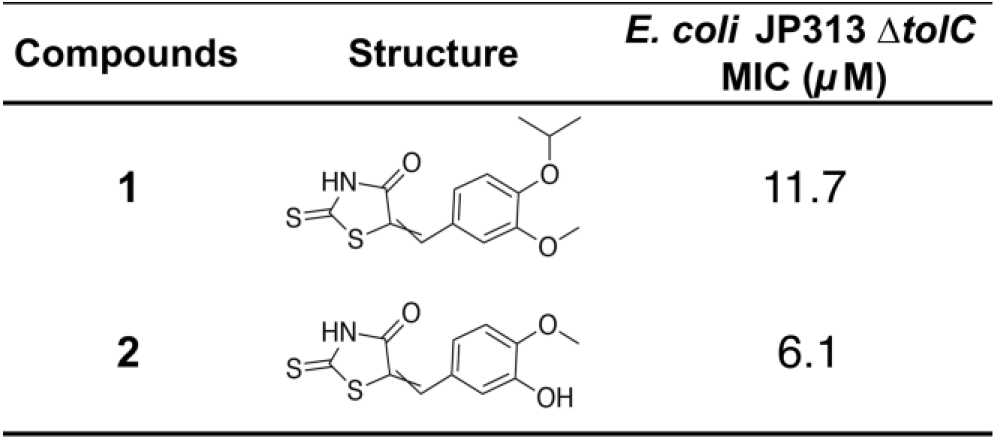
Structures and MICs of PAINS compounds 1 and 2.

| Compounds | Structure | *E. coli* JP313 $\Delta tolC$<br>MIC ($\mu M$) |
| --- | --- | --- |
| 1 |  | 11.7 |
| 2 |  | 6.1 |

**Figure 1.**
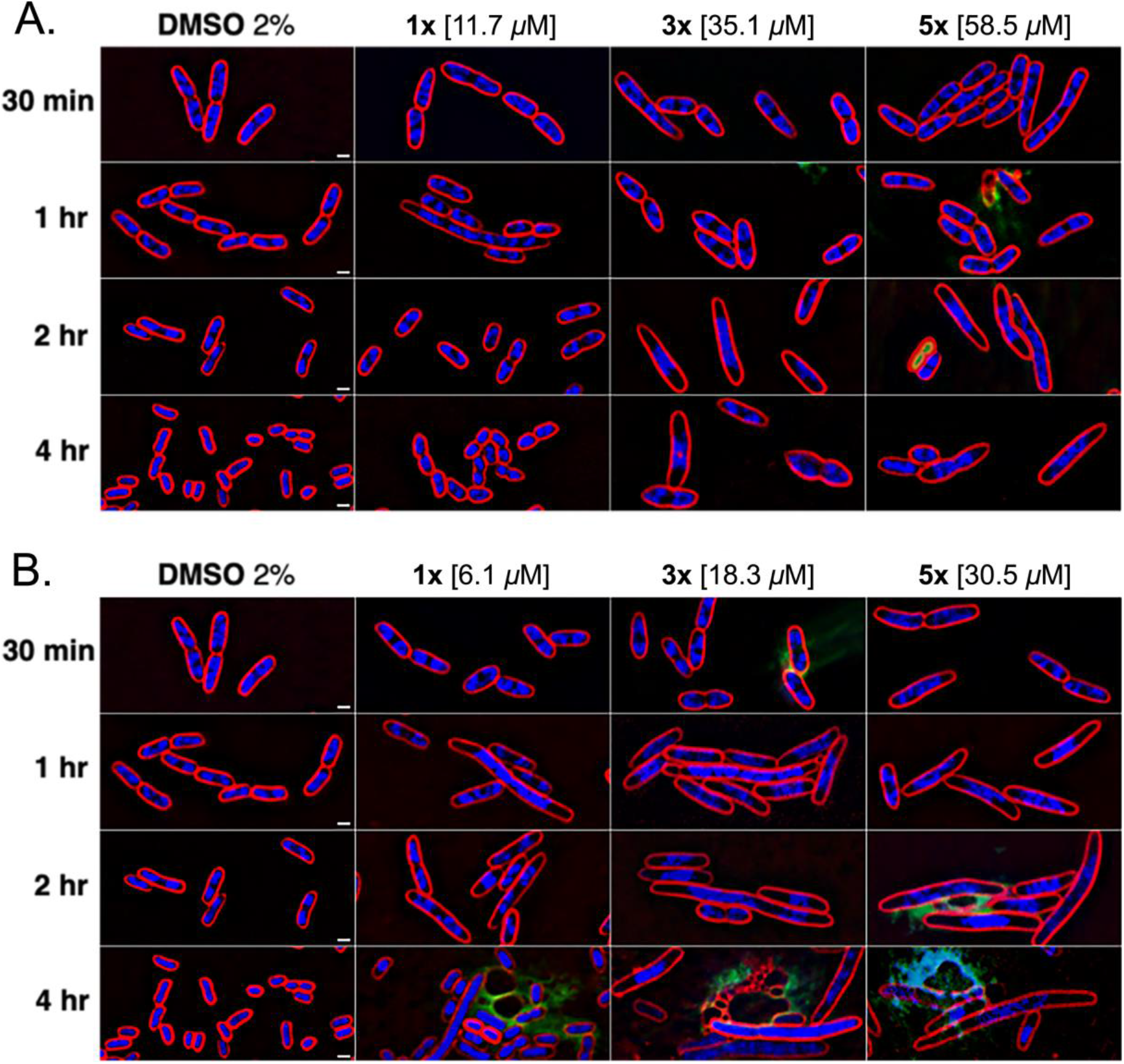
Cytological profiles of E. coli JP313 ΔtolC treated two PAINS. Cells were exposed to **(A)** compound 1 or **(B)** compound 2 at 1X, 3X, and 5X MIC for 30 minutes, one hour, two hours, and four hours. Images were taken after staining the cells with FM4-64 (red), DAPI (blue), and SYTOX-green (green). The scale bar represents one micron.

Our viability experiments further characterized the antibacterial effects of compounds 1 and 2 against *E. coli ΔtolC*. Compound 1 appeared largely bacteriostatic, for even when cells were treated at 5X MIC, viability never decreased considerably (Figure 2A). Compound 2, on the other hand, caused nearly a 100-fold reduction in the number of viable *E. coli ΔtolC* cells after treatment at 5X MIC (Figure 2B). These data are consistent with our BCP observations because, while both compounds inhibited DNA replication, compound 2 appeared to inhibit cell wall biogenesis to a much greater degree and caused significant cell lysis.

**Figure 2.**
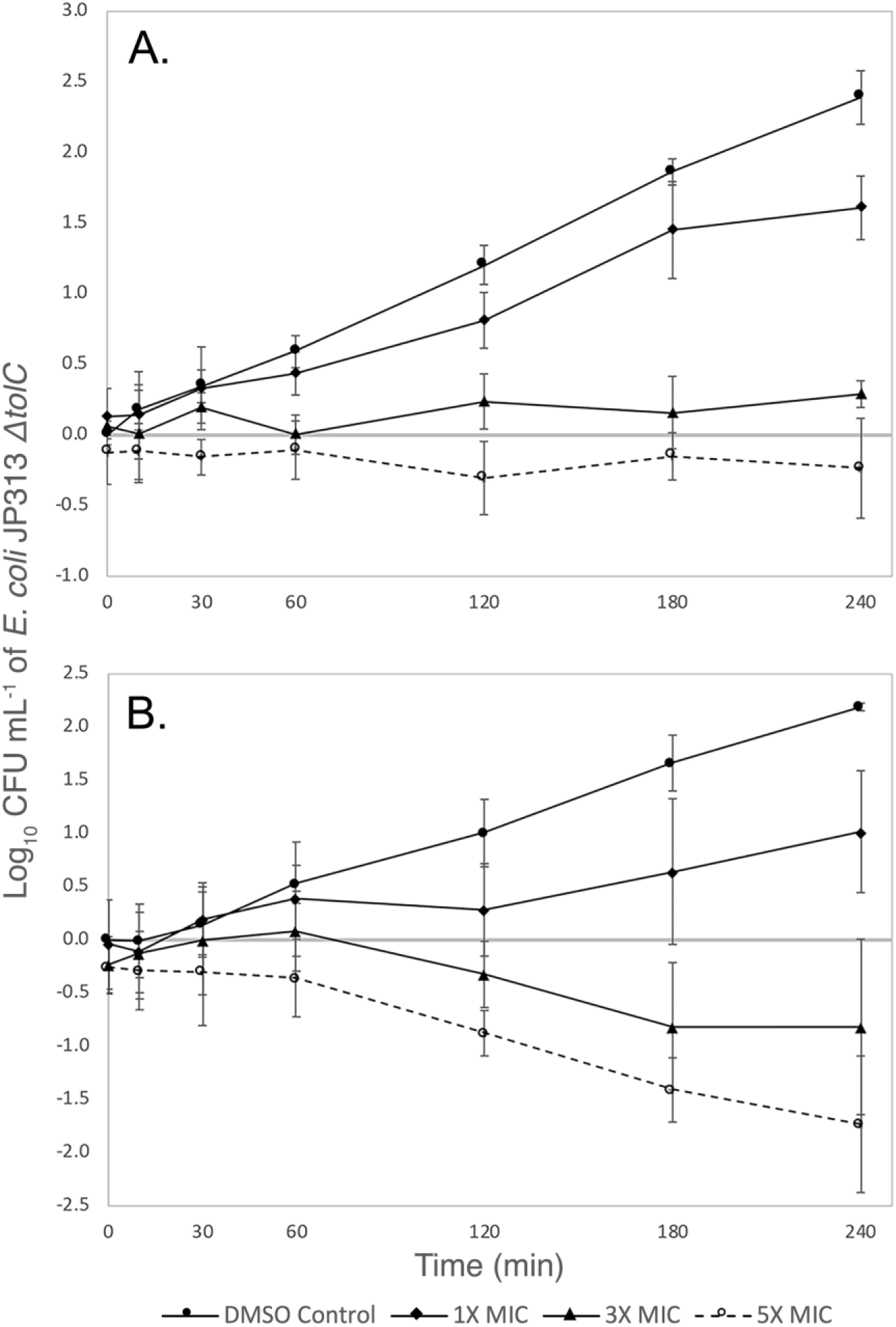
Viability of E. coli JP313 ΔtolC treated with two PAINS. Cells were treated with varying concentrations of **(A)** compound 1 or **(B)** compound 2. CFU/mL was measured at 30 minutes, one hour, two hours, three hours, and four hours. Standard error values were calculated from three independent trials.

### Identifying TMPK as a cellular target by isolating resistant mutations and performing *in silico* docking

Though we suspected that compounds 1 and 2 inhibited the DNA replication pathway, we did not know which protein within the pathway was inhibited. In order to identify a potential enzymatic target, we passaged *E. coli ΔtolC* serially in the presence of increasing concentrations of both compounds and obtained one resistant mutant for each compound. The MIC of compound 1 against its corresponding resistant mutant (mutant 1) was six times higher than the MIC against the *E. coli ΔtolC* parent strain, and mutant 1 was also resistant (~2.8-fold) to compound 2 (Table 2). Mutant 2 (from compound 2) was roughly 10-fold resistant to both compounds. Whole genome sequencing revealed that both resitant mutants contained missense mutations in the *tmk* gene, which codes for TMPK (Figure S2). In mutant 1, proline replaced glutamine at residue 45, and in mutant 2, threonine replaced alanine at residue 69. Mutant 1 contained an additional mutation in the *dksA* gene, and mutant 2 contained an additional mutation in the *yjeA* gene. However, because *tmk* was the only gene that was mutated in both strains and because cross-resistance was conferred, we suspected that these mutated *tmk* alleles were likely responsible for conferring resistance to the compounds. Therefore, we moved the *tmk* allele from the stronger resistant mutant into a clean strain background via phage P1 transduction. In isolation, the *tmk(A69T)* allele conferred 6-fold resistance to compound 1 and 2.4-fold resistance to compound 2 (Table 2). These results suggest that TMPK is a cellular target of both compounds *in vivo.*

**Table 2.**
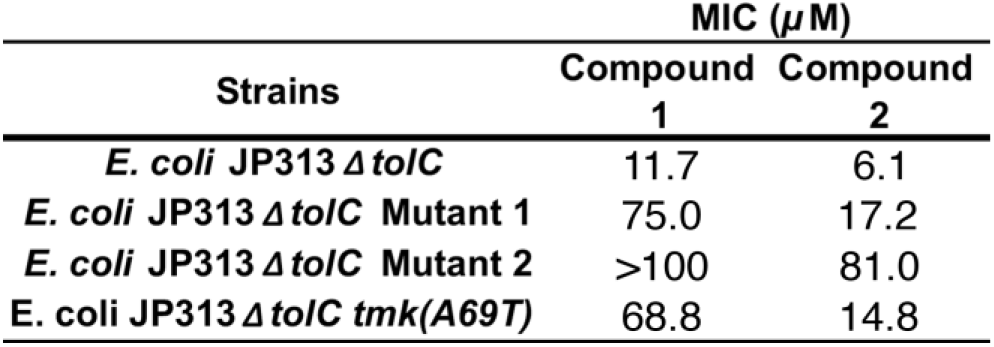
MICs of compounds 1 and 2 against resistant mutants.

| Strains | MIC (μM) | |
| --- | --- | --- |
|  | Compound 1 | Compound 2 |
| *E. coli* JP313 ΔtolC | 11.7 | 6.1 |
| *E. coli* JP313 ΔtolC Mutant 1 | 75.0 | 17.2 |
| *E. coli* JP313 ΔtolC Mutant 2 | >100 | 81.0 |
| *E. coli* JP313 ΔtolC *tmk*(A69T) | 68.8 | 14.8 |

In order to probe the potential molecular mechanism of our compounds, we used computational modeling to dock compounds 1 and 2 to TMPK (Figure 3). This analysis revealed binding sites for compounds 1 and 2 that overlap considerably within the TMPK active site. The high-probability binding orientation of our inhibitors closely mimicked that of the substrate (thymidine monophosphate) and known analog inhibitors.^26, 27^ It is likely that by binding at this location, compounds 1 and 2 compete with thymidine monophosphate binding and block the enzyme’s catalytic activity. Neither of the resistant mutations that we selected were located in the TMPK active site, and it remains unclear how these mutations confer resistance to compounds 1 and 2.

**Figure 3.**
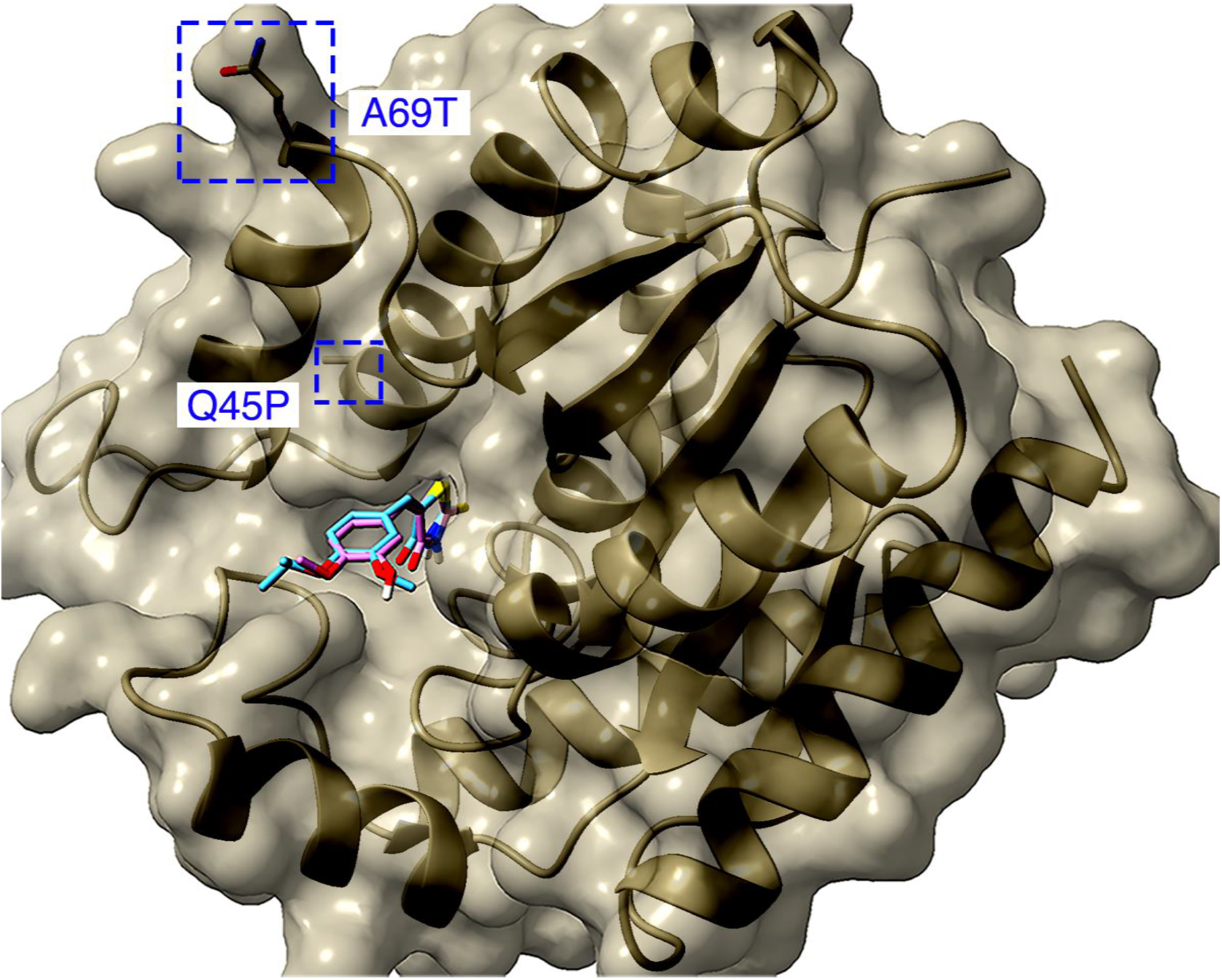
Compounds 1 and 2 docked to the active site of TMPK. Compound 1 (cyan) and compound 2 (magenta) are shown overlapping. Mutated residues conferring resistance are labeled in blue. Heteroatoms are colored such that sulfur is yellow, oxygen is red, and nitrogen is blue.

### Screening Analogs of Compounds 1 and 2

In order to identify specific inhibitors of TMPK, we screened 29 structural analogs of compounds 1 and 2 for their antibacterial activity and MOA against *E. coli ΔtolC* (Figure S3). Of these analogs, seven showed antibacterial activity with MICs less than 50 *μ*M (Table 3, Table S1). Compound 27 was the most potent, with an MIC of 1.9 *μ*M. BCP was used to determine whether these analogs inhibit DNA replication specifically or have other mechanisms of action (Figure 4). Many of the compounds induced filamentation and chromosomal replication defects, indicating that they inhibit DNA replication (Figure 4), but most of these molecules also inhibited other pathways such as cell wall biogenesis, which was evidenced by cell lysis. Compound 13 induced membrane permeability and cell lysis with no sign of inhibited DNA replication. This compound had a relatively high MIC compared to most other active molecules. In addition to nonspecific inhibitors, we identified three molecules (compounds 9, 11, and 27) that selectively inhibited DNA replication. The molecule with perhaps the most promise as a TMPK inhibitor was compound 27, as it displayed the lowest MIC and a clear cytological profile indicating high specificity for the DNA replication pathway.

**Table 3.**
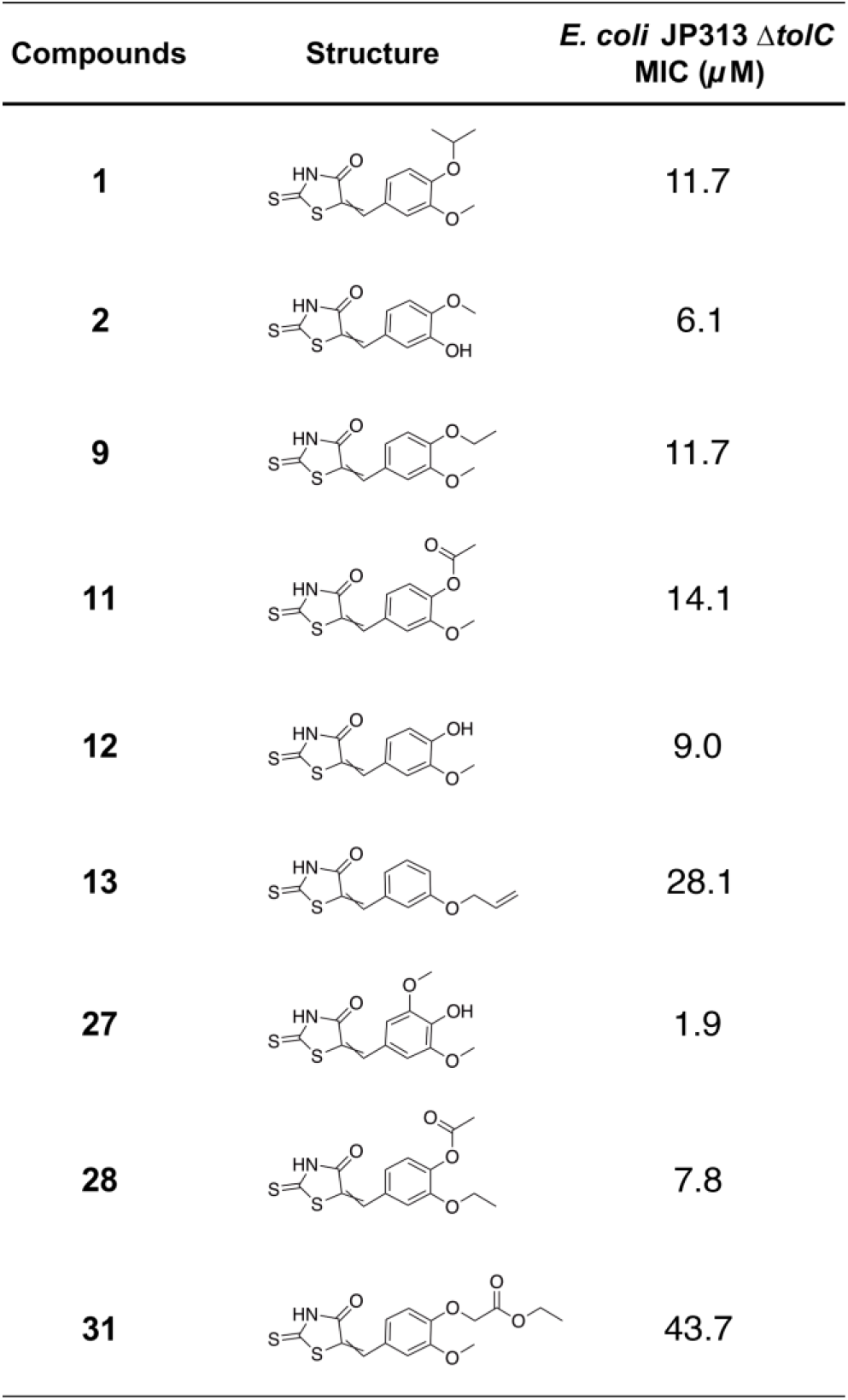
Structures and MICs of PAINS analogs.

| Compounds | Structure | *E. coli* JP313 $\Delta tolC$<br>MIC ($\mu M$) |
| --- | --- | --- |
| 1 |  | 11.7 |
| 2 |  | 6.1 |
| 9 |  | 11.7 |
| 11 |  | 14.1 |
| 12 |  | 9.0 |
| 13 |  | 28.1 |
| 27 |  | 1.9 |
| 28 |  | 7.8 |
| 31 |  | 43.7 |

**Figure 4.**
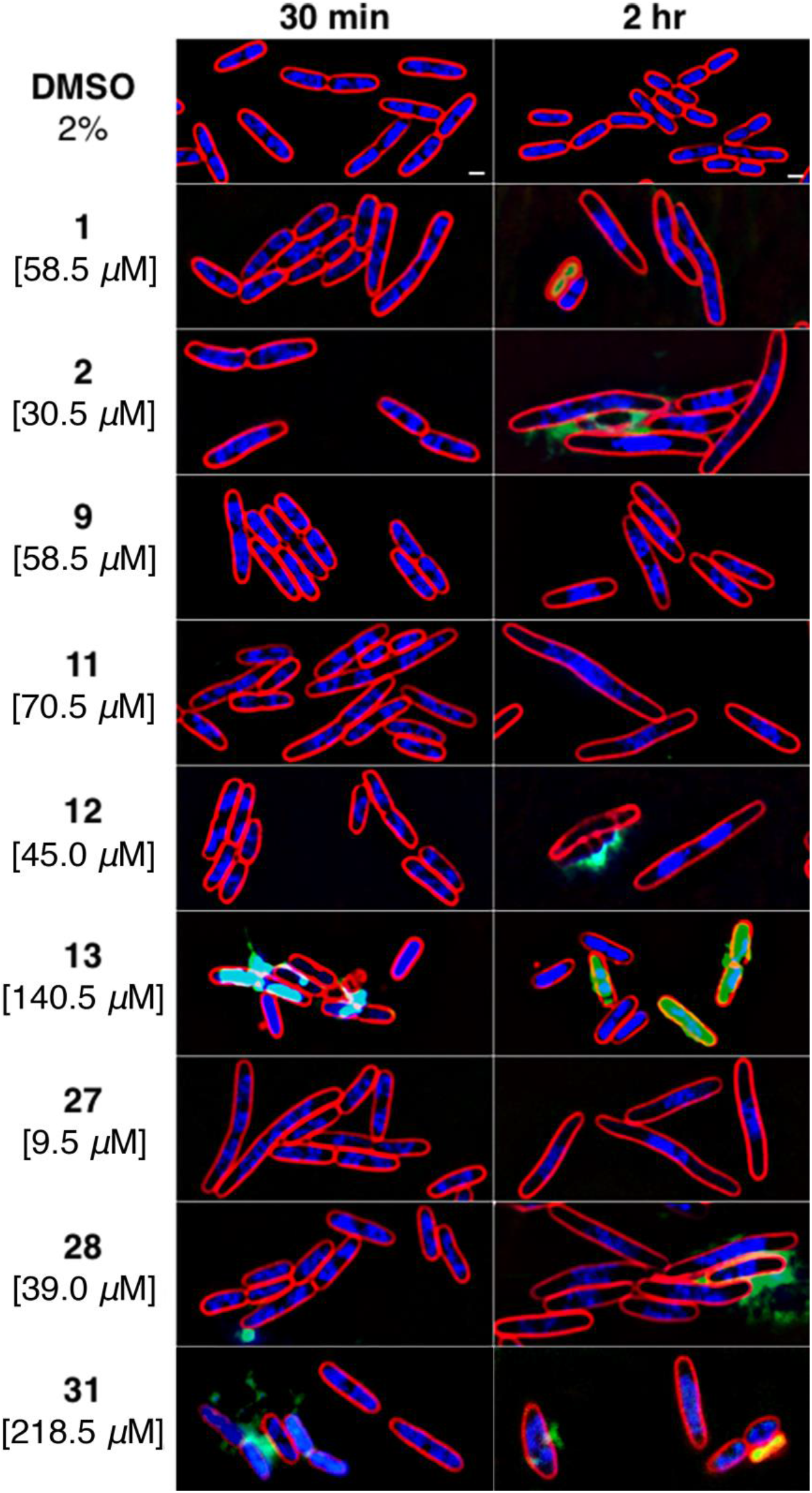
Cytological profiles of E. coli JP313 ΔtolC treated with PAINS analogs. E. coli JP313 ΔtolC cells were treated with 2% DMSO as a control. Analogs were administered at 5X MIC (concentrations shown on the left), and images were taken after 30 minutes and two hours. The scale bar represents one micron.

### Testing mutant and plasmid-overexpressing strains for resistance to analogs

To provide evidence that TMPK is the molecular target of some of our PAINS analogs, we tested the activity of these molecules against the *tmk(A69T)* mutant strain. This mutant exhibited some degree of resistance to all of the analogs (Table 4). The MIC of compound 27 was most drastically increased (10-fold) in the *E. coli* strain containing *tmk(A69T)*, further suggesting that this analog specifically inhibits thymidylate kinase *in vivo*. Conversely, those analogs that appeared not to inhibit DNA replication as strongly as compound 27 showed only a modest increase in MIC against the *tmk(A69T)* strain. For example, the MICs of compounds 13 and 31 were only approximately 2-fold higher against this strain compared to the strain with wild-type *tmk*.

**Table 4.** MICs of PAINS analogs against E. coli ΔtolC tmk(A69T) and plasmid overexpressing strains as well as IC50 values for the inhibition of E. coli TMPK in vitro.

| Compounds | Allele-conferred Resistance (μM) | | | Plasmid Overexpression (μM) | | | | *In vitro* Inhibition (μM) |
| --- | --- | --- | --- | --- | --- | --- | --- | --- |
| | *E. coli* $\Delta tolC$ *tmk*<sup>+</sup> | *E. coli* $\Delta tolC$ *tmk*(A69T) | Conferred Fold Resistance | Vector Control | Plasmid + *tmk* | Plasmid + *tmk*(Q45P) | Plasmid + *tmk*(A69T) | *E. coli* TMPK $IC_{50}$ |
| 1 | 11.7 | 68.8 | 5.9 | 15.5 | 62 | 62 | 62 | 0.39 |
| 2 | 6.1 | 14.8 | 2.4 | 7.8 | 31 | 62 | 62 | 2.50 |
| 8<sup>a</sup> | >100 | - | - | - | - | - | - | 1.20 |
| 9 | 11.7 | 43.8 | 3.7 | 15.5 | 62 | 62 | 62 | 0.41 |
| 11 | 14.1 | 37.5 | 2.7 | 15.5 | 62 | >62 | >62 | 0.48 |
| 12 | 9.0 | 20.3 | 2.3 | 7.8 | 31 | >62 | >62 | 0.50 |
| 13 | 28.1 | 56.3 | 2.0 | 31.3 | 62.5 | 62.5 | 62.5 | 1.00 |
| 14<sup>a</sup> | >100 | - | - | - | - | - | - | 4.20 |
| 16<sup>a</sup> | >100 | - | - | - | - | - | - | 5.60 |
| 27 | 1.9 | 18.8 | 10 | 3.9 | 15.5 | 15.5 | 15.5 | 0.25 |
| 28 | 7.8 | 19.4 | 2.5 | 7.8 | 31.3 | 64 | 64 | 0.47 |
| 31 | 43.7 | 93.8 | 2.1 | 31.3 | 125 | 125 | 125 | 0.68 |

<sup>a</sup> compound was inactive *in vivo*

To provide additional evidence that thymidylate kinase might be the target of this family of compounds, we tested whether TMPK overexpression conferred resistance. To accomplish this, we cloned the wild-type *tmk* gene as well as the *tmk(A69T)* and *tmk(Q45P)* resistant alleles on multi-copy number plasmids and measured the MICs of all nine of our active compounds against *E. coli ΔtolC* transformed with these plasmids. Overexpression of the wild-type or mutant *tmk* genes resulted in at least a 4-fold increase in the MICs of all compounds with BCP phenotypes characteristic of DNA replication inhibitors (Table 4). In the cases of compounds 2, 12 and 28, an even greater increase in MIC (up to 8-fold) was observed when either *tmk(A69T)* or *tmk(Q45P)* was overexpressed. By contrast, the MIC of compound 13, which had no obvious effect on DNA replication *in vivo,* was increased only 2-fold upon overexpression of *tmk* alleles.

### PAINS analogs inhibit thymidylate kinase *in vitro*

Because our genetic and cell biology studies suggested that thymidylate kinase is a target of this family of molecules, we determined whether or not these compounds inhibited the purified *E. coli* thymidylate kinase *in vitro*. We obtained IC_50_ values for all nine active compounds and also for three compounds with no activity against *E. coli ΔtolC* (compounds 8, 14, and 16) (Table 4). The inactive compounds studied were chosen on the basis of their structural diversity. The IC_50_ values for all 12 compounds ranged from 250 nm (compound 27) to 5.6 *μ*M (compound 16) (Table 4). As might be expected, the three compounds for which MICs could not be measured had some of the highest IC_50_ values and yielded poor dose-response curves (Figure S4). Compound 27, which is the most potent and specific inhibitor *in vivo*, displayed the lowest IC_50_. Despite this, our data seemed to demonstrate only a weak correlation between compound MIC and IC_50_. This correlation was primarily affected by our limited dataset as well as the unexpectedly high IC_50_ value associated with compound 2, one of the strongest inhibitors *in vivo*. Any inconsistencies between MICs and IC_50_ values can be explained by a number of potential contributing factors. First, many of these compounds have multiple cellular targets that contribute to their *in vivo* activity. Second, the cell envelope is a significant barrier to antibiotic infiltration, and different structural features of our antibiotic analogs could affect their ability to enter the cell.^28^ Finally, some of these molecules might behave as PAINS and yield misleading data when studied using *in vitro* assays such as this one.

Here we present a collection of genetic, cell biological, and biochemical data that demonstrate that many of the molecules within a particular rhodanine-containing chemical series inhibit cellular TMPK. Though some of these molecules have additional targets *in vivo,* three compounds appeared to specifically target TMPK. Compound 27, stands out among this group because in addition to its specificity (eliciting clear DNA replication defects *in vivo*), it had the most potent IC_50_ against the purified protein *in vitro* and inhibited the growth of *E. coli ΔtolC* with the lowest MIC. While it may be the case that PAINS cannot be developed as antibiotics, this work nonetheless high-lights the utility of combining an *in vivo* MOA assay with medicinal chemistry to study the antibacterial mechanisms of molecules.

## Materials and Methods

### Construction of *E. coli* JP313 *ΔtolC*

The *ΔtolC* mutation is derived from strain PB3^29^ and was introduced into strain JP313^30^ by P1 transduction. JP313 was transduced to tetracyline resistance with a lysate of strain CAG18475 (*metC162*::Tn*10*), and the methionine requirement of the transductants was confirmed. This strain was then transduced to prototrophy with a lysate of PB3, and these transductants were screened on MacConkey agar for the presence of the *ΔtolC* mutation. Transductants that acquired *ΔtolC* were unable to grow on this medium owing to their sensitivity to bile salts such as deoxycholate,^31^ and these arose at about 6%, in keeping with previous linkage data.^29^ PB3 and CAG18475 were obtained from the *Coli* Genetic Stock Center at Yale University.

### Synthetic compound library screen

A synthetic small molecule library consisting of ~1,800 compounds was obtained from the ChemBridge EXPRESS-pick library stock collection and screened in 96-well plates at 100 *μ*g/mL in LB to identify compounds active against *E. coli* JP313 *ΔtolC*.

### Determining MICs

MICs were determined using the broth microdilution method performed in triplicate. All compounds used in this study were purchased from ChemBridge and solubilized in dimethyl sulfoxide (DMSO). Concentrated stocks of each compound were prepared at 20 mM and stored at −80°C. A 1:100 dilution of an overnight culture of *E. coli* JP313 *ΔtolC* was prepared in LB liquid media and grown at 30°C to an OD_600_ between 0.2 and 0.4. With the exception of a media control column, 1 *μ*L of cells diluted to an OD_600_ of 0.05 was added to a 96-well plate containing 2-fold serial dilutions of eight different starting concentrations of each compound in 100 *μ*L LB. An initial cell density count was determined using a plate reader prior to incubation at 30°C for 24 hours while shaking. After 24 hours, the optical density was determined, and OD_600_ readings were corrected by subtracting the initial reading at T^0^ from the final OD_600_ reading at T^24^. Readings of 0.5 and below constituted inhibition.

### Bacterial cytological profiling (BCP)

High-resolution fluorescence microscopy and BCP were performed as previously described by Nonejuie et. al.^19^ Briefly, overnight cultures of *E. coli* JP313 *ΔtolC* were diluted 1:100 in LB and incubated with rolling at 30°C until they reached an OD_600_ between 0.15-0.17. 300 *μ*L of cells was treated with compounds prepared to the desired test concentration and incubated while rolling at 30°C. Microscopy images were taken after 30 minutes, one hour, two hours, and/or four hours. Prior to applying the treated culture to an agarose pad for imaging, the cells were stained with FM4-64, DAPI, and SYTOX-green as previously described.^19^

### Cell viability

Viability assays were performed in triplicate for *E. coli* JP313 *ΔtolC* treated with compounds 1 and 2. Cells were plated for colony counting at the same time points used for BCP imaging. The plates were incubated at room temperature overnight, then individual colonies were counted. The number of colony-forming units per mL was calculated for each treatment condition. These values were then normalized using the T^0^ measurement of the untreated cells and subsequently log_10_ transformed. The resulting viability measurements were averaged across three independent trials and plotted with standard errors.

### Resistant mutant selection

Mutants resistant to compounds 1 and 2 were obtained via serial passaging. A 1:500 dilution of an overnight culture of *E. coli* JP313 *ΔtolC* was prepared in 6 mL LB and incubated at 30°C with rolling until the cells reached an OD_600_ of 0.2-0.25. In a small glass culture tube, 1 *μ*L of cells was added to 1 mL of LB containing compound at a concentration of 0.5X MIC. The cultures were incubated at 30°C for 1 day while rolling. If growth occurred at the starting concentration, the culture was diluted 1000-fold using LB containing a slightly higher concentration of compound. Serial passaging was performed until the highest concentration of each compound was reached, at which point colony purification was performed on plates containing the compound.

### Genomic DNA extraction and quantification

Genomic DNA was extracted using a protocol adapted for use with Qiagen’s DNeasy® Blood & Tissue Kit. The gDNA concentration was quantified using a Thermo ScientificT™ NanoDrop™ One Microvolume UV-Vis Spectrophotometer. gDNA was stored at −20°C.

### Genome sequencing, assembly, and variant analyses

The genome sequences of two *E. coli* JP313 *ΔtolC* mutants were generated using the Illumina MiSeq sequencing platform with a V2 500 paired-end read kit at the La Jolla Institute for Allergy and Immunology Research, San Diego, CA. All sequence processing and analyses were performed in Geneious Prime. Two FastQ files containing forward and reverse sequence read lists were generated for each genome. The files were imported into Geneious and simultaneously paired (input of expected insert size of 500 bp), resulting in a single paired read list. The sequences were trimmed using the BBDuk v38.37 plugin. BBDuk identifies and trims adapters based on presets for Illumina adapters and paired read overhangs, trims the end of sequences based on quality (Q score), and discards short reads and their associated pair mate. Adapter sequences trimmed using presets for Illumina adapters were trimmed from the right end of the sequence with the Kmer length set at 27 with a maximum substitution allotment of 1. A minimum overlap value of 25 was set for adapter trimming based on paired read overhangs. Low quality sequences were trimmed on both ends with the minimum quality score set at 30, and reads shorter than 30 bp were discarded. After trimming, the paired-end reads were merged into a single read with BBMerge. The resulting merged and unmerged sequences were mapped to the previously sequenced and annotated genome of the *E. coli* JP313 *ΔtolC* parent strain using the Geneious assembler set to medium sensitivity, and to find structural variants, short insertions, and deletions of any size. Further variation analysis was performed with the resulting alignment using the Geneious SNP finder, set to use the bacterial genetic code and the following parameters: a minimum variant frequency of 0.25, maximum variant P-value of 6, minimum strand-bias P-value of 5 when exceeding 65% bias, and to calculate an approximate P-value for identified variants. Visual inspection of these resulting data led to the identification of mutations present in each genome.

### Primer design and polymerase chain reaction (PCR)

Thymidylate kinase DNA templates (~1,099 bp) were amplified from *E. coli* JP313 *ΔtolC* strains using a Q5 high-fidelity DNA PCR (New England Biolabs). The primers used for these reactions were “EM029-TMKF1” (5’-ATGGCAAACTTTCTCGTG-3’) and “EM030-TMKR1” (5’-GGTGTAGTAATCGGGATG-3’). Each PCR mixture (50 *μ*L) contained 100 ng of template genomic DNA, 500 pmol of each primer, and 200 *μ*M dNTPs. The thermocycling conditions were 30 seconds of initial denaturation at 98°C followed by 30 cycles of denaturation (98°C, 10 seconds), annealing (61°C, 15 seconds), and extension (72°C, 2 minutes). A final extension was performed at 72°C for 5 minutes. PCR products were purified with the oligonu-cleotide cleanup protocol as described in the Monarch PCR & DNA Cleanup Kit (5 *μ*g) user manual (NEB #T1030). Purified PCR products were sequenced using Sanger methods by Eton Biosciences (https://www.etonbio.com/) and trimmed for quality prior to analysis.

### *E. coli* genome manipulation by P1 virulent transduction

P1 transductions of the *tmk(A69T)* allele were carried out according to the protocol described by Miller and Miller.^32^ The *tmk(A69T)* mutation was backcrossed into *E. coli* JP313 *ΔtolC* by cotransduction with the closely linked *fhuE* gene. A P1*vir* lysate of strain JW1088-5 [Δ*fhuE764*::*kan*] was used to transduce the original *tmk(A69T)* mutant to kanamycin resistance, and transductants were screened for retention of resistance to compound 2. A P1*vir* lysate of one such transductant was then used to transduce *E. coli* JP313 *ΔtolC* to kanamycin resistance. Colonies that were also resistant to compound 2, having acquired the *tmk(A69T)* mutation by cotransduction, represented the desired backcross. JW1088-5 was obtained from the Coli Genetic Stock Center at Yale University.

### TMK expression in *E. coli* JP313 *ΔtolC*

Three full length *E. coli* K-12 MG1655 thymidylate kinase alleles (WT, Q45P, and A69T) were amplified and cloned into the plasmid expression vector pRSFDuet-1 (GenScript). Expression is under control of the T7 lac promoter. Plasmids were transformed into *E. coli* JP313 *ΔtolC* to give the following strains: JP313 *ΔtolC*-pRSFDuet, JP313 *ΔtolC*-pRSFDuet-*tmk*^+^, JP313 *ΔtolC*-pRSFDuet-*tmk*-134, and JP313 *ΔtolC*-pRSFDuet-*tmk*-205.

These strains were grown in LB media in the presence of kanamycin (30 *μ*g/mL) to an OD_600_ of 0.1−0.2, at which point they were treated with the compounds. The assay was performed with eight different concentrations of each compound, serially diluted 2-fold, in 96-well plates in 100 *μ*L of LB with kanamycin 30 *μ*g/mL. Cell densities were determined using a plate reader before and after a 24-hour incubation at 30°C with shaking. Final OD_600_ values were determined by subtracting the values from the initial reading at T^0^. MICs were determined as the lowest concentration of compound resulting in an OD_600_ of 0.5 or below, indicating growth inhibition.

### Molecular docking

The protein docking was performed using Autodock Vina (v1.1.2). The crystal structure of Escherichia coli thymidylate kinase (TMPK) was obtained in complex with P1-(5’-adenosyl)-P5-(5’-thymidyl)pentaphosphate and P1-(5’-adenosyl)P5-[5’-(3’-azido-3’-deoxythymidine)] pentaphosphate from Protein Data Bank (PDB code: 4TMK, http://www.rcsb.org/).^26^ Water molecules, cofactors, and the originally docked ligand were removed while polar hydrogens and partial charges were added with Autodock Tools (v1.5.6), which is available as a part of MGLtools v1.5.6 through The Scripps Research Institute (http://mgltools.scripps.edu/downloads). Docking was carried out following the standard procedures for Autodock Vina and the resulting docking configuration was viewed and illustrated with Chimera X (https://www.rbvi.ucsf.edu/chimerax/).^33^

### *E. coli* TMPK IC_50_ determination

Inhibition of *E. coli* TMPK catalysis was measured in the direction of ATP synthesis using the ATPlite luminescence assay (PerkinElmer cat. no. #6016739) in white 384-well plates (Corning cat. no. #3825). The assay buffer consisted of 50 mM HEPES (pH 8.0), 25 mM sodium acetate, 10 mM MgCl_2_, 5 mM dithiothreitol, 0.01% Triton X-100, and 0.5 mM EDTA. The thymidine 5’-diphosphate (TDP) and adenosine 5’-diphosphate (ADP) concentrations were 65 mM and 10 mM, respectively. The TMPK concentration was 400 pM. Test compounds stock solutions were prepared at 10 mM in DMSO. Serial 2-fold dilutions were prepared in buffer supplemented with DMSO such that the final DMSO concentration in the assay was constant at 0.33% and the highest compound concentration tested was 33 mM. A replicate plate was prepared with no TMPK to serve as the 100% inhibition control and to allow correction for signal suppression by the test compounds, according to the method described in Shapiro et al (2009).^34^ The assay volume during the 1-hour TMPK reaction was 10 mL. The reactions were quenched by addition of 5 mL of ATPlite reagent. Background luminescence was measured in wells containing only buffer and ATPlite reagent. Luminescence was measured, after 5 min of incubation in the dark, using a Pherastar FS plate reader (BMG Labtech), with 1-s integration/well.

The IC_50_ values were calculated as follows. The average luminescence of the background wells was subtracted from the luminescence of all the other wells. The correction for signal suppression was made using the data from the plate with no TMPK. The average uninhibited control luminescence (MAX) was measured in the wells containing TMPK but no inhibitor. The average fully-inhibited control (MIN) was measured in the wells containing no inhibitor or TMPK. The percent inhibition at each inhibitor concentration was calculated for each compound using the equation: % inhibition = 100[1-(X-MIN)/(MAX-MIN)], where X is the measurement in the well with TMPK and the specific concentration of inhibitor. The IC_50_ was calculated by nonlinear regression from the % inhibition data using the Excel add-on XLfit (ID Business Solutions Ltd., U.K.), equation 205, in which the minimum and maximum % inhibition are fixed at 0 and 100, respectively: % inhibition = 100[I]^n^/(IC_50_^n^ + [I]^n^), where [I] is the inhibitor concentration and n is the Hill slope.

## ASSOCIATED CONTENT

### Supporting Information

The Supporting Information is available in the file:

TMK_Supporting_Information (PDF)

## AUTHOR INFORMATION

**Authors:**

**Elizabeth T. Montaño**‡ **-**Division of Biological Sciences, University of California, San Diego, La Jolla, California, USA.

**Jason F. Nideffer**‡ **-**Division of Biological Sciences, University of California, San Diego, La Jolla, California, USA.

**Joseph Sugie -**Division of Biological Sciences, University of California, San Diego, La Jolla, California, USA.

**Eray Enustun -**Division of Biological Sciences, University of California, San Diego, La Jolla, California, USA.

**Adam B. Shapiro -**Entasis Therapeutics, Gatehouse Park BioHub, Waltham, Massachusetts, USA.

**Hannah Tsunemoto -**Division of Biological Sciences, University of California, San Diego, La Jolla, California, USA.

**Alan I. Derman -**Division of Biological Sciences, University of California, San Diego, La Jolla, California, USA.

**Kit Pogliano -**Division of Biological Sciences, University of California, San Diego, La Jolla, California, USA.

ETM, JFN, JS, EE, ABS, HT, and AID conducted experiments and analyzed data. The manuscript was written by ETM, JFN, AID, KP, and JP. All authors have read and approved the manuscript.

## ACKNOWLEDGMENT

These studies were supported by the National Institute of Health (R01AI177112). KP and JP have an equity interest in Linnaeus Bioscience Incorporated and receive consulting income from the company. The terms of this arrangement have been reviewed and approved by the University of California, San Diego in accordance with its conflict-of-interest policies.

## ABBREVIATIONS

BCP: bacterial cytological profiling
DMSO: dimethyl sulfoxide
IC_50_: half maximal inhibitory concentration
MDR: multi-drug resistant
MIC: minimum inhibitory concentration
MOA: mechanism of action
PAINS: panassay interference compounds
SAR: structure-activity relationship
TMPK: thymidylate kinase

